# LXR agonist Prevents Peripheral Neuropathy and modifies PNS immune cells in Aged Mice

**DOI:** 10.1101/2021.09.22.461263

**Authors:** Chaitanya K. Gavini, Nadia Elshareif, Anand V. Germanwala, Gregory Aubert, Nigel A. Calcutt, Virginie Mansuy-Aubert

**Affiliations:** Cell and Molecular Physiology, Stritch School of Medicine, Loyola University Chicago, Maywood, Illinois, USA 60153; Department of Neurological Surgery, Loyola University Medical Center, Maywood, Illinois, USA 60153; Departement of Internal Medicine, Division of Cardiology, Loyola University Medical Center, Maywood, Illinois, USA 60153; Department of Pathology, University of California San Diego, La Jolla, California, USA 92093

**Keywords:** Liver X receptors, GW3965, peripheral neuropathy, aging

## Abstract

Peripheral neuropathy is a common and progressive disorder in the elderly that interferes with daily activities and increases the risk of injury. It is of importance to find efficient treatments to treat or delay this age-related neurodegeneration. We previously demonstrated that activation of the cholesterol sensor Liver X receptor (LXR) with the potent agonist GW3965, alleviates pain in a diet-induced obesity model. Because cholesterol had also been linked to neuropathy during aging, we sought to test whether LXR activation may improve neuropathy and pain in aged mice by treating 21-month-old mice for 3 months with GW3965. Treatment resulted in a significant increase in nerve fibers of the sub-basal plexus, accompanied by a change in polarization, metabolism, and cholesterol content of macrophages in the sciatic nerve. These results suggest that activation of the LXR may block the progression of neuropathy associated with aging by modifying nerve-immune cell cholesterol, thereby providing new pathways to target in efforts to delay neuropathy during aging.

## Introduction

Aging-related peripheral neuropathy and neuropathic pain contributes significantly to decreased quality of life in the elderly. Surveys report a prevalence of neuropathy and neuropathic pain of between 20-58% in people aged 60 and above, with prevalence directly proportional to age [1–4]. With age, the peripheral nervous system (PNS) undergoes pathophysiological changes that are readily observed in people suffering with neuropathy [5]. In humans, aging leads to atrophy of large myelinated fibers and thinning of the myelin sheath [6, 7]. Similar to humans, rodent models of peripheral neuropathic pain may develop pathological sensitivity to light touch (allodynia) [8, 9].

Many forms of disease-initiated peripheral neuropathy share a common pathogenic mechanism that involves modifications to neuro-immune interactions mediated by inflammatory signals [10]. Recent studies have demonstrated that an increase in the number of macrophages in the dorsal root ganglia (DRG) and the sciatic nerve (SN) alters pain perception [11–13]. Macrophages and neutrophils were shown to be pivotal actors in demyelinating diseases such as Wallerian degeneration or after nerve transection [14, 15]. Macrophages helps with nerve regrowth after phagocytosis of myelin debris [14]. During nerve transection, macrophages form a bridge to facilitate migration of cells such as endothelial cells and Schwann cells [16, 17]. These functions contribute to nerve repair; however, macrophages can also have detrimental function. For example in the Guillain-Barre syndrome, macrophages can penetrate myelinating fibers at the nodes of Ranvier that leads to axonal damage [18]. In model of Charcot-Marie-Tooth (CMT), endoneurial macrophages are activated by Schwann cells and damage nerve fibers [19]. Martini et al. showed that macrophage silencing by blocking a cytokine receptor reduced endoneurial foamy macrophages and significantly improves age-related degenerative changes in peripheral nerves of 24 months-old mice [7].

Circulating cholesterol and cholesterol pathways are linked to development and progression of neuropathy and studies have identified lipid-filled macrophages in the nerves of 24 months old mice [20–22]. The function of lipid-loaded macrophages is still obscure in the PNS but a pool of them are likely a consequence of clearing of membrane debris. The clearance of this foamy macrophages - extensively studied in atherosclerosis-is fundamental and dysfunction of lipid efflux from macrophages leads to inflammatory pathology [23]. Liver X receptors (LXR) are ligand-activated nuclear receptors that bind metabolites of cholesterol [24, 25], regulate macrophage lipid efflux [26, 27] and are involved in progression of early type II diabetic neuropathy [28]. Oxysterols derived from phagocytosis of apoptotic or necrotic cells activate the LXR pathway in naive macrophages, upregulating genes involved in cholesterol efflux such as Abca1 and ATP binding cassette A1 [29]. Overexpression of ABCA1 induces expression of the anti-inflammatory cytokines and other markers of M2 macrophage [30–32]. The role of LXR in immune cells located in the nerve system had not been investigated. In this study, we therefore treated aged mice with the selective LXR agonist GW3965 and profiled immune cell changes within the DRG and SN. We observed enhanced nerve regeneration and amelioration of neuropathic pain in aged mice treated with GW3965 that may be the consequence of changes in nerve macrophage cholesterol efflux.

## Materials and methods

### Mice

All studies were conducted in accordance to recommendations in the Guide for the Care and Use of Laboratory Animals of the National Institutes of Health and the approval of the Loyola University Chicago Institutional Animal Care and Use Committee. C57BL/6J (#000664) were obtained from Jackson laboratory (Maine, USA). All mice were housed under a 12:12 h light/dark cycle. Mice had free access to food (Teklad LM-485: Envigo, Indiana, USA) and water. All studies were performed using male mice with experimenter blinded to treatment using the ARRIVE guidelines.

### Human subjects

Informed consent was obtained from all human subjects prior to sural nerve sample collection in strict accordance with the rules and guidelines stipulated by the Loyola University Chicago Internal Review Board (IRB). All experimental protocols were approved by the Loyola University Chicago IRB (protocol # 210567020519).

### In vivo agonist treatment

Aged mice were treated with vehicle or LXR agonist (GW3965; 25mg/kg BW) (Axon Medchem, Virginia, USA) by i.p. twice weekly for 12 weeks, starting at 21 months of age. Tissues were rapidly dissected and processed or frozen in liquid nitrogen before analysis.

### Mechanical Sensitivity

Mice were investigated for mechanical allodynia using phasic stimulation with von Frey filaments as described [9]. Briefly, mice were acclimated to the testing chambers for 40 minutes and were then subjected to stimulations with 6 calibrated von Frey filaments (0.16; 0.4; 1; 2; 4; 6; 8 g: North Coast Medical, California, USA). Filaments were applied 6 times for 1 sec at 1 sec intervals with a 5 min break between each series of stimulations. Response frequency for each filament was recorded and 50% threshold calculated. A single trained investigator took all baseline and experimental measurements for these series of experiments while remaining blinded to the treatment groups. Mice were evaluated in a quiet room, at a constant temperature of 24°C and acclimated to the von Frey chambers for at least 40 minutes, but not restrained in the chamber any longer than necessary to minimize stress and discomfort-induced behavioral variations. Allodynia was characterized as an intense paw withdrawal or licking of the stimulated hind paw [33].

### Thermal nociception

Mice were investigated for heat nociception using Plantar Test Apparatus (Hargreaves Method: IITC Life Science, California USA) [33]. Briefly, after acclimation to testing chambers, tests were performed on the hind paw plantar surface using a focused, radiant heat light source with a built-in timer displaying reaction time in seconds. A cutoff time of 20 seconds was set to avoid tissue damage.

### Nerve Fiber Densities

Foot pads were collected from hind paws and transferred to Zamboni’s fixative for 4 to 6 h on ice. Samples were processed for immunohistochemistry, nerve fibers identified by immunostaining with anti-PGP9.5 antibody (#7863-0504, AbD Serotec) and intra-epidermal quantified as described in detail elsewhere [33].

### Flow Cytometry

Cells dissociated from the DRG and SN were stained with fluorescently conjugated antibodies (CD45, CD11b, I-A/I-E, F4/80, CD206) as previously described [34]. Cells that were positive for both CD45 and CD11b were sorted from other dissociated cells for further analysis. Flow cytometry data were acquired using a BD FACS Aria III and data analyzed using FlowJo (Treestar).

### Quantitative PCR

mRNA was extracted from sorted CD45+/CD11b+ cells using Arcturus PicoPure RNA Isolation Kit (ThermoFisher) before generating cDNA using High Capacity cDNA Reverse Transcription Kit (ThermoFisher). For all genes of interest, qPCR was performed using Sybr green-based assay (Roche, Indiana, USA) using IDT primers (IDT technologies, Iowa, USA). 18s was used to normalize data and quantification was done using ΔΔCT method with vehicle treated group’s mean value set at 100%.

### Cholesterol Assay

Cholesterol content in sorted CD45+/Cd11b+ cells was assessed using the Amplex Red Cholesterol Kit (ThermoFisher) following manufacturer’s instructions. Data are normalized to protein content with the vehicle treated group’s mean value set at 100%.

### Immunofluorescence

Bone derived macrophages were obtained from C57BL/6J mice as previously described [35]. Briefly, femurs and tibia were obtained from mice after euthanasia and marrow flushed using RPMI-1640 supplemented with 10% FBS, antibiotics, and glutamine. The fluid containing dissociated marrow cells was passed through a cell strainer. Cells were plated onto coverslips coated with poly-l-lysine in a 12-well plate and allowed to attach overnight in a humidified incubator with 5% CO_2_ at 37°C. The following day, coverslips were washed 3 times with warm phosphate buffered saline (PBS) to remove non-adherent cells before the media was replaced with RPMI-1640 supplemented with 10% FBS, antibiotics, glutamine, and 10% L929 conditioned media to differentiate the cells. Once fully differentiated (~ day 6), cells were treated with either vehicle or 1 μM GW3965 for 24 hours, then co-incubated with 10μg/mL DiI-oxLDL (ThermoFisher) for another 24 hours. Cells were then washed with ice cold PBS and fixed with 4% PFA for 15 minutes. After washing with PBS, coverslips were co-stained with DAPI and mounted for imaging. Images were captured using Olympus IX80 Inverted Microscope equipped with an X-Cite 120Q fluorescent light source (Lumen Dynamics) and a CoolSNAP HQ2 CD camera (Photometrics). Image processing and quantification was performed using CellSens (Olympus Corporation, Waltham, Massachusetts) and ImageJ software.

### Oxygen consumption assay

A Seahorse XFe96 Analyzer (Agilent) was used to measure oxygen consumption rate (OCR) [36–38] of macrophages (seeding density 100,000 cells/well) treated with oxLDL and GW3965 as described above. Briefly, 1hr before the assay, culture media was aspirated from the cells in the plate and Seahorse XF assay media supplemented with 20mM glucose and 2mM glutamine was applied and equilibrated in a CO_2_ free incubator. Oligomycin (2μM, final) (inhibits complex V of the electron transport chain (ETC)), carbonyl cyanide-4-(trifluoromethoxy)phenylhydrazone (FCCP, uncoupling agent) (2μM, final), antimycin A and rotenone (0.75μM, final) (inhibit complexes I and III, inhibiting ETC activity) were sequentially injected into the system to induce changes in ETC activity. OCR values were normalized to total protein in individual wells.

### Quantification and statistical analysis

All data are represented as Mean±S.E.M. Analysis and graphing were done using GraphPad Prism 9.1.2. For single group comparisons, either a 1- or 2-tailed t-test was used as appropriate. Multiple comparisons were performed using one-way ANOVA. The number of experiments/replicates and mice for each experiment are described in figure legends.

## Results

### Aged mice developed features of neuropathic pain and small fiber neuropathy

We used the von Frey and Hargreaves tests to longitudinally evaluate sensitivity of 6-month, 12-month, and 21-month-old mice. During the von Frey test, we observed a significant decrease in the 50% response threshold of mice starting at 12 months of age and progressing at 21 months of age (Figure1A) suggesting an increased sensitivity to innocuous stimuli with age. Mice also developed sensitivity to heat, with significant hyperalgesia detected in 21-month-old mice compared to 12-month-old mice (Figure1B). Thus, age significantly affects escape responses to both mechanical and thermal stimuli.

**Figure1:**
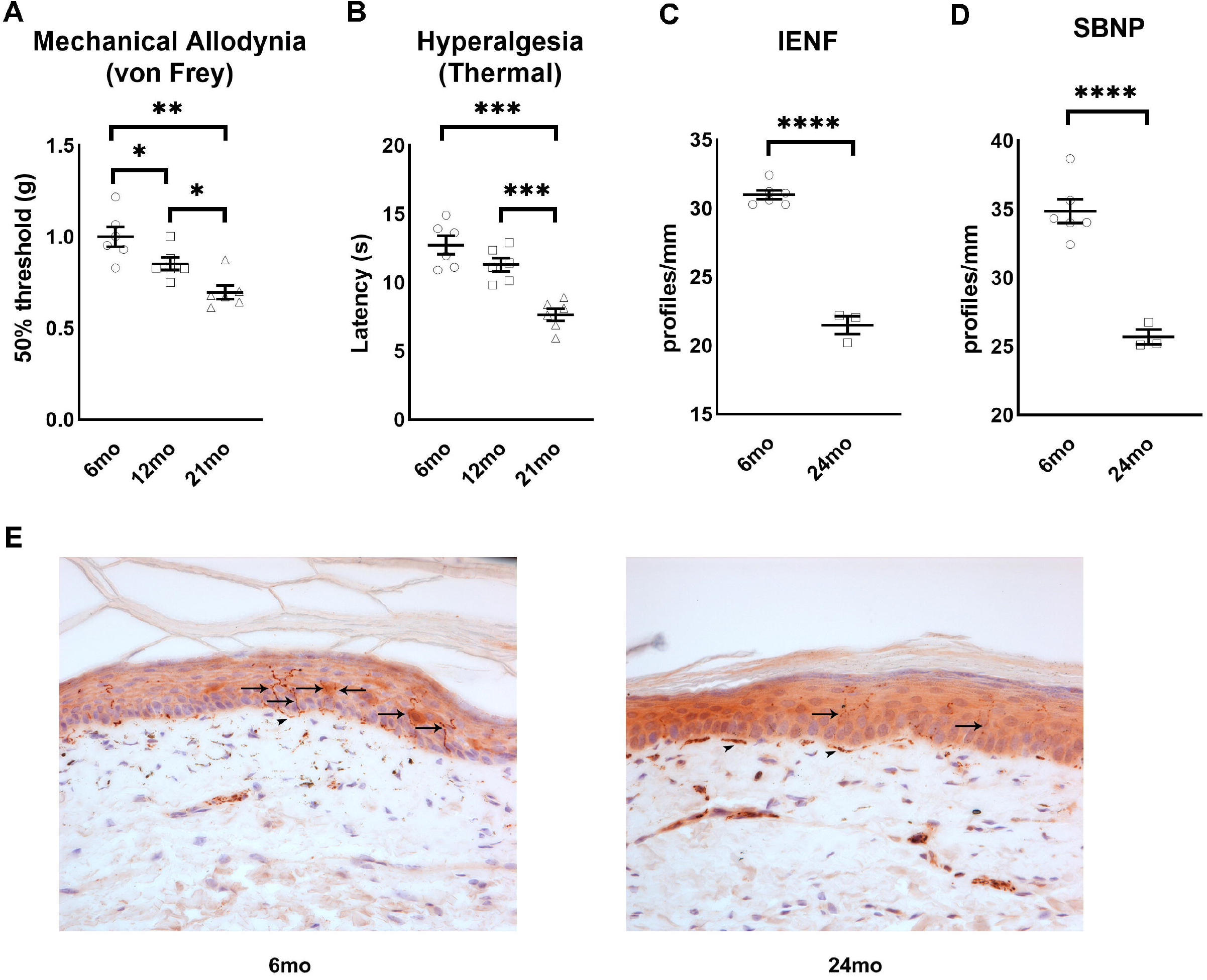
Aging leads to loss of distal nerve and neuropathy. A) Mechanical allodynia in 6-month, 12-month, and 21-month-old mice (n=6/group). B) Thermal hyperalgesia in 6-month, 12-month, and 21-month-old mice (n=6/group). Distal nerve quantification in 6-month and 24-month-old mice, IENF (C) and SBNP (D) (n=3-5/group). E) Representative images of paw plantar skin of 6-month and 24-month-old mice used for IENF quantification. PGP9.5 positive nerve fibers stain brown. Black arrows indicate intra-epidermal nerve fibers and arrowheads indicate dermal nerves. All images were obtained at 40x magnification. All data are Mean±SEM. *p<0.05, **p<0.005, ***p<0.0005, ****p<0.00005.

We then quantified sensory nerve density in the epidermis (intraepidermal nerve fibers; IENF and sub-basal nerve plexus; SBNP). We observed a significant decrease in both IENF (Figure1C, E) and SBNP (Figure1D, E) density in 24-month-old mice compared to 6-month-old mice demonstrating that aging led to distal sensory neuropathy.

### Age-related accumulation of macrophages in the dorsal root ganglia and sciatic nerve

Previous studies in mice have identified macrophages as contributors to assorted peripheral neuropathies [39]. In order to assess the immune cell population, we performed flow cytometry of cells derived from the DRG and SN and used specific cell surface markers to identify macrophage numbers and phenotypes (Figure 2A). In line with current literature, we observed a significant temporal increase in the percentage of cells expressing the pan-macrophage markers CD45+F4/80+ in both the DRG and SN, with the highest increase occurring between 12 and 24 months of age (Figures 2B, C). We also observed a temporal increase in percentage of cells identified as pro-inflammatory M1 macrophages (CD45+IAIE+) in the DRG and SN (Figures 2B, C) and a significant increase in the percentage of the anti-inflammatory M2 macrophage (CD45+CD206+) population in both DRG and SN of 12-month-old mice compared to 6-month-old mice (Figures 2B, C). However, the proportion of M2 macrophages significantly decreased in the DRG and SN of 24-month-old mice (Figure2B, C).

**Figure2:**
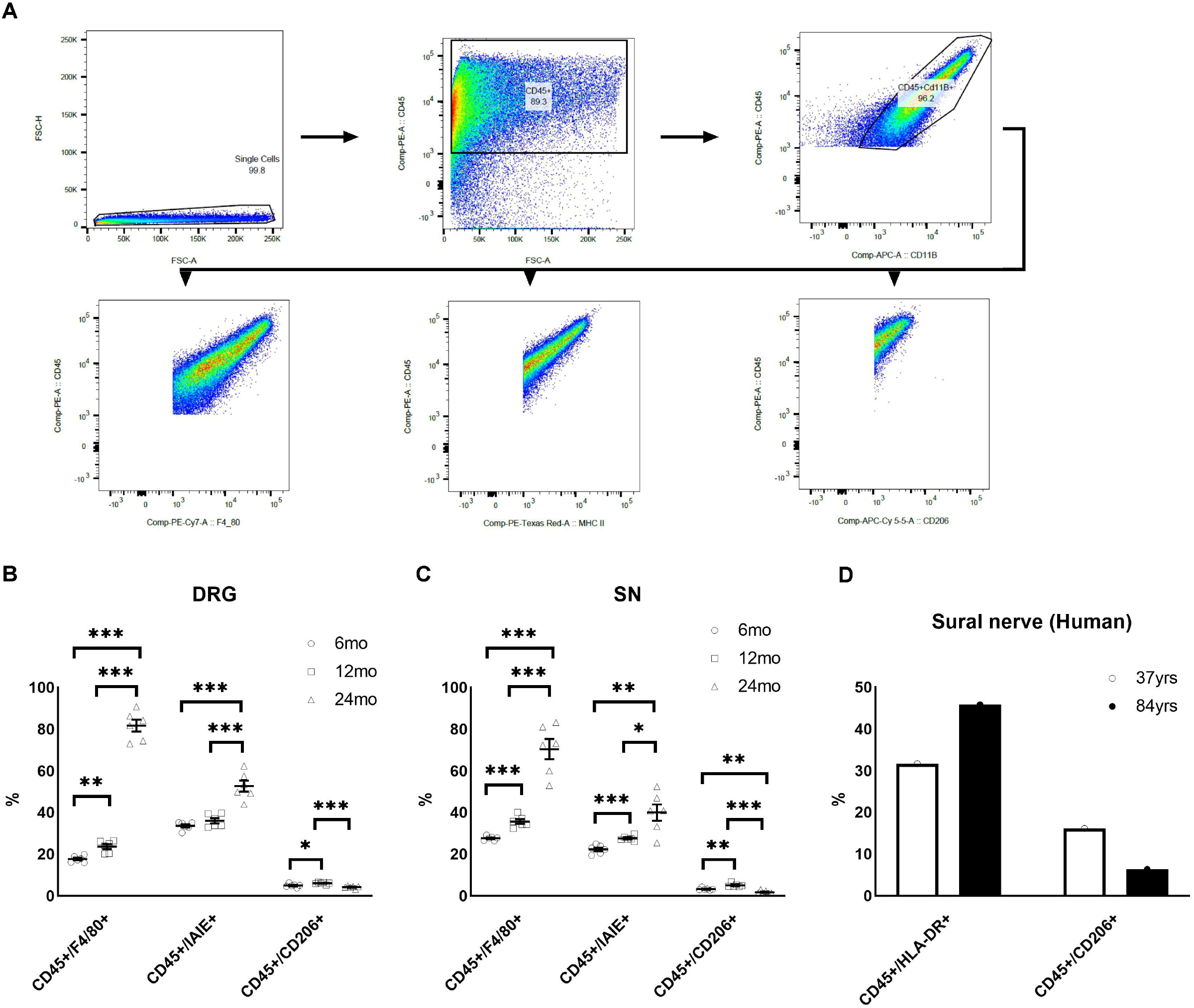
Activated macrophage numbers and their phenotype change with age. A) Gating strategy used for sorting and quantifying macrophage population. B) Percentage macrophage population and phenotypes in the DRG of 6-month, 12-month, and 21-month-old mice (n=6/group). C) Percentage macrophage population and phenotypes in the SN of 6-month, 12-month, and 21-month-old mice (n=6/group). D) Percentage of M1 (CD45+/HLA-DR+) and M2 (CD45+/CD206+) macrophages in human sural nerve biopsies. All data are Mean±SEM. *p<0.05, **p<0.005, ***p<0.0005.

Comparison of sural nerve biopsies from two individuals (subject 1: 84 years and subject 2: 37 years) demonstrated a considerably lower percentage of anti-inflammatory M2 macrophages in older individual compared to the younger individual (Figure 2D). Similar to our mice data, we also observed an increase in M1 macrophages in older individual compared to younger individual (Figure2D). These data demonstrate that phenotype of activated macrophages changes with age.

### LXRs agonist delays progression of age-associated neuropathic pain and neuropathy

We next assessed whether activation of LXRs using its potent and selective agonist GW3965 could change the progression of age-related neuropathic pain and neuropathy. 21-month-old mice were injected with LXRs agonist, GW3965 (25mg/kg body weight), or vehicle for 12 weeks [9]. Compared to vehicle, treatment with GW3965 prevented the development of mechanical hypersensitivity and thermal hyperalgesia over time with a significant difference observed after 6 weeks of treatment for mechanical allodynia and after 9 weeks of treatment for thermal sensitivity (Figures 3A, B, C, and D). We then evaluated the effect of GW3965 on IENF and SBNP density in the paw of these mice. Compared to vehicle treated mice, we observed an increasing trend in IENF of 24-month-old mice treated with GW3965 (Figures 3E, G) and a significant increase in SBNP density in 24-month-old mice treated with GW3965 (Figures 3F, G). This data suggests that LXRs activation either attenuated distal degeneration or stimulated nerve regeneration and demonstrate that in vivo LXRs activation using GW3965 can improve indices of age-related neuropathic pain and sensory neuropathy.

**Figure3:**
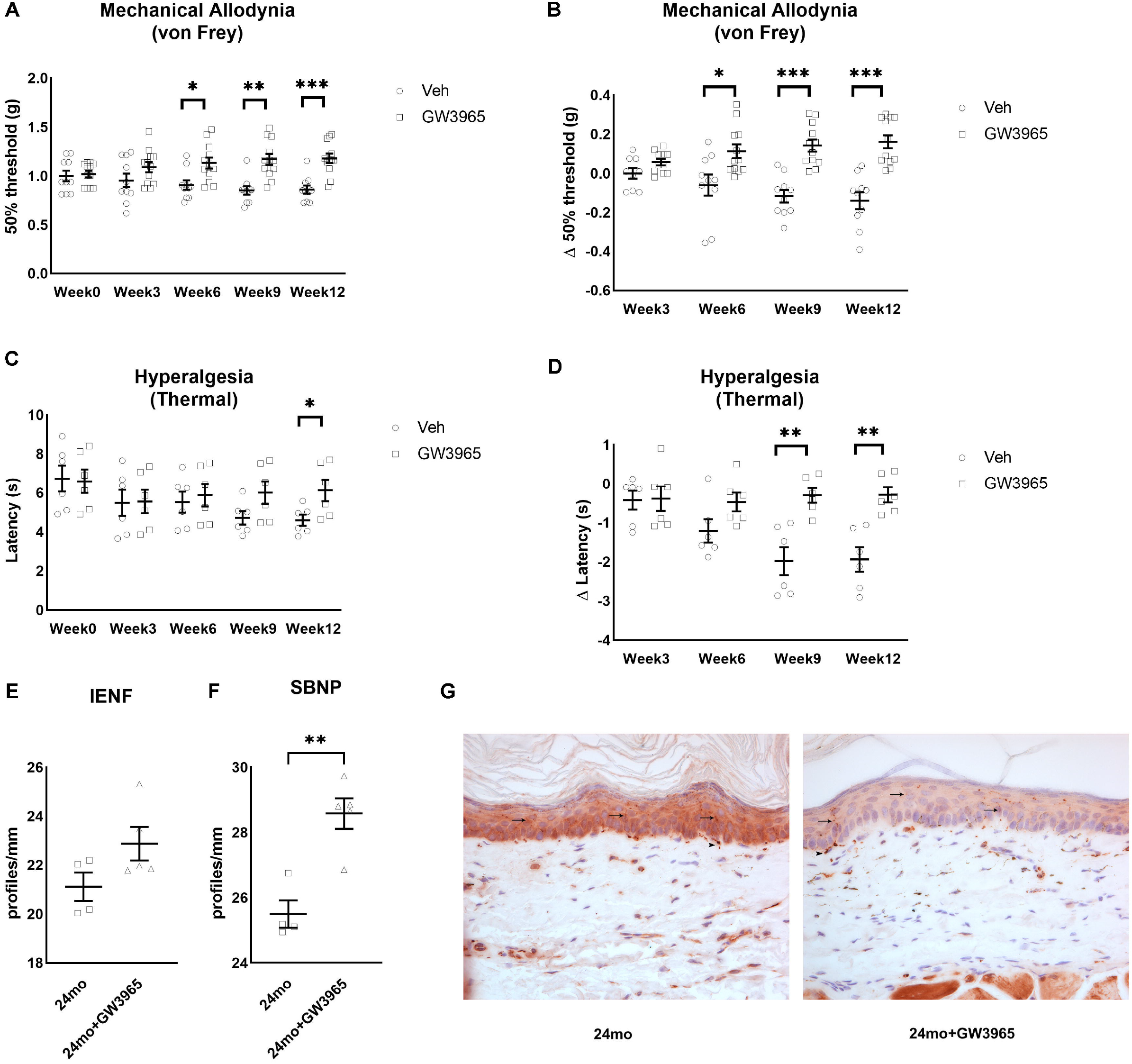
LXRs activation improves indices of age-related neuropathic pain. A) Mechanical allodynia in old mice treated with and without LXRs agonist GW3965 for 12 weeks (n=10/group). B) Change from baseline in mechanical allodynia in old mice treated with and without LXRs agonist GW3965 (n=10/group). C) Thermal hyperalgesia in old mice treated with and without LXRs agonist GW3965 for 12 weeks (n=6/group). D) Change from baseline in thermal hyperalgesia in old mice treated with and without LXRs agonist GW3965 (n=6/group). Distal nerve quantification in 24-month-old mice treated with and without LXRs agonist GW3965, IENF (E) and SBNP (F) (n=4-5/group). G) Representative images of paw plantar skin of 24-month-old mice treated with and without LXRs agonist GW3965 used for IENF quantification. PGP9.5 positive nerve fibers stain brown. Black arrows indicate intra-epidermal nerve fibers and arrowheads indicate dermal nerves. All images were obtained at 40x magnification. All data are Mean±SEM. *p<0.05, **p<0.005, ***p<0.0005.

### Effect of LXRs agonist treatment on macrophage polarization in the DRG and SN

To assess if LXR activation also modifies PNS immune cell number and phenotype we quantified the immune cell population in the DRG and SN using flow cytometry. Although there was not a significant change in the DRG (Figure 4A), we did observe a significant increase in percentage of both pro-inflammatory (CD45+IAIE+) M1 macrophages and anti-inflammatory (CD45+CD206+) M2 macrophages in the SN of mice treated with GW3965 (Figure 4B).

**Figure4:**
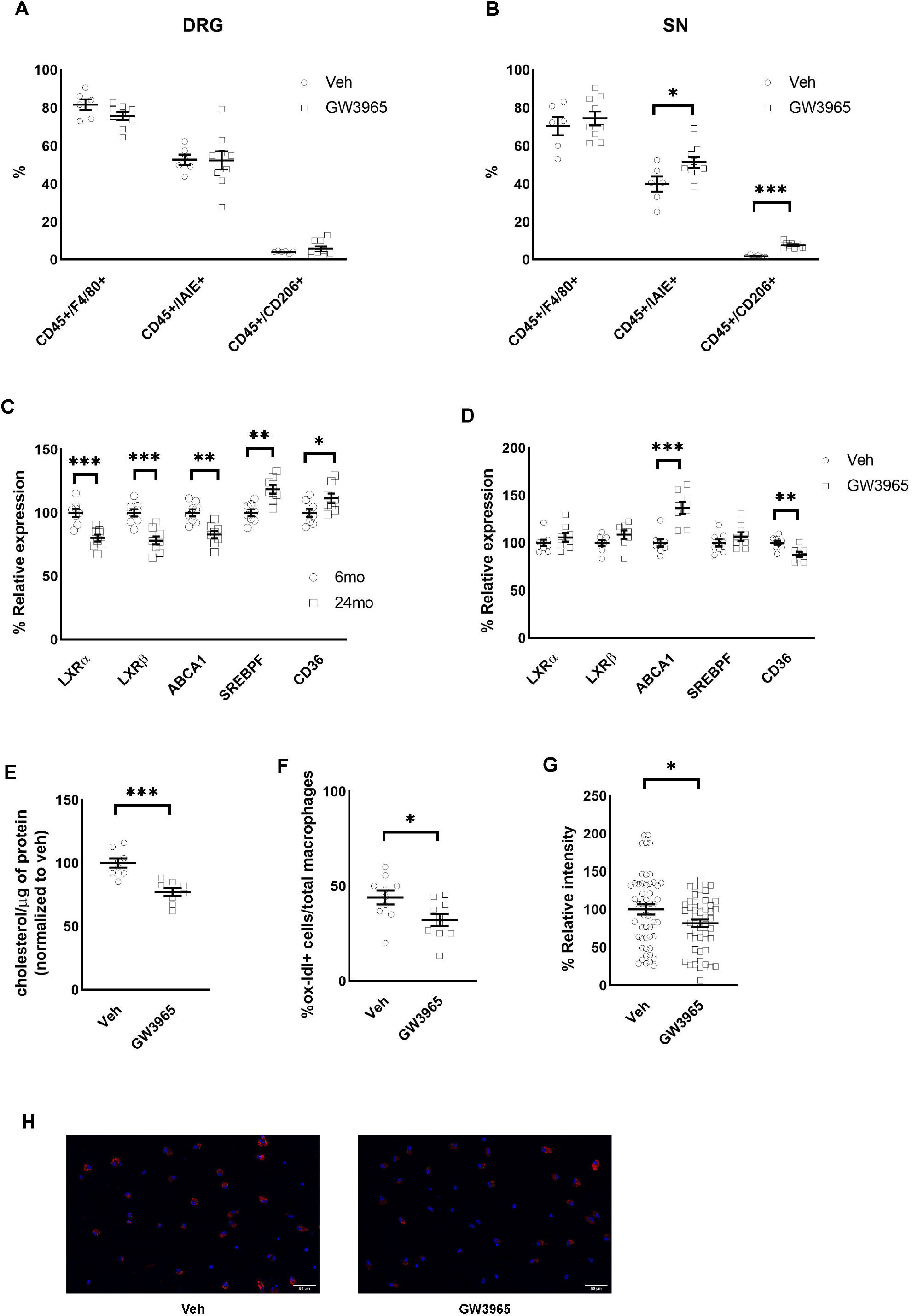
Activation of LXRs increases lipid efflux. A) Percentage macrophage population and phenotypes in the DRG of old mice treated with and without LXRs agonist GW3965 (n=6-9/group). B) Percentage macrophage population and phenotypes in the SN of old mice treated with and without LXRs agonist GW3965 (n=6-9/group). C) mRNA expression of LXRs target genes in the sorted CD45+Cd11b+ cells from the SN of young and old mice (n=8/group). D) mRNA expression of LXRs target genes in the sorted CD45+Cd11b+ cells from the SN of old mice treated with and without LXRs agonist GW3965 (n=8/group). E) Cholesterol content in the sorted CD45+Cd11b+ cells from the SN of old mice treated with and without LXRs agonist GW3965 (n=8/group). F) Percentage oxldl positive cells from macrophages treated with and without LXRs agonist GW3965 (n= 3 experiments in triplicate). G) oxldl relative intensity in oxldl positive cells from macrophages treated with and without LXRs agonist GW3965 (n= 50 cells/group). H) Representative images of oxldl positive cells from macrophages treated with and without LXRs agonist GW3965. All data are Mean±SEM. *p<0.05, **p<0.005, ***p<0.0005.

Our data show age-related decrease in gene expression of efflux mediators e.g. abca1 (Figure4C). LXRs play an important role in regulating macrophage cholesterol efflux in other tissues [26, 27] but their role in regulation of cholesterol in macrophages of the nerves had never been evaluated to our knowledge. We therefore investigated whether restoring the efflux capacity of macrophages in aged mice could improve their functions. We sorted CD45+CD11b+ cells from the DRG and SN and assessed a variety of genes associated with lipid homeostasis including production of lipid de novo and lipid efflux : Abca1, Srebpf (Sterol regulatory element-binding transcription factor 1), and Cd36 (cluster of differentiation 36, scavenger receptor) in LXRs treated mice. We found a significant increase in expression of Abca1 and a significant decrease in expression of CD36 (Figure 4D). We also found significantly decreased cholesterol content in sorted cells from mice treated with GW3965 compared to vehicle treated mice (Figure 4E). These data suggest that LXR activation in vivo regulates the balance lipid uptake/efflux within immune cells of peripheral nerves. As sorted cells (CD45+CD11b+) represent a non-specific myeloid cell population, we extended our studies to bone marrow derived macrophages. Differentiated macrophages derived from bone marrow were treated with oxidized low-density lipoproteins (oxldl) to induce lipid-loaded cells [40, 41], and then treated with GW3965. Consistent with current literature, we found a significantly lower percentage of cells labeled with oxldl in GW3965 treated group (Figure 4F, H). Within the cells that were positive for oxldl, GW3965 treated cells had significantly lower content of oxldl compared to vehicle treated cells (Figure 4G, H).

To test if LXRs activation affected macrophage metabolism, we measured oxygen consumption rate (OCR) in differentiated macrophages derived from bone marrow treated with oxldl with and without GW3965 (Figure 5A). Compared to vehicle, LXRs agonist treatment significantly increased basal respiration (Figure 6B) and basal glycolytic activity (basal ECAR, Figure 6C). LXRs activation also significantly increased maximal respiration (Figure 6D), proton leak (Figure 6E), and non-mitochondrial oxygen consumption (Figure 6F), without having a significant effect on spare respiratory capacity, coupling efficiency, or ATP-linked oxygen consumption (not shown). These data suggest that LXRs activation change the metabolism of macrophages exhibiting energetic phenotype. Altogether, our findings indicate that increased lipid efflux via activation of LXRs and ABCA1 modify macrophage in the PNS of aged mice.

**Figure5:**
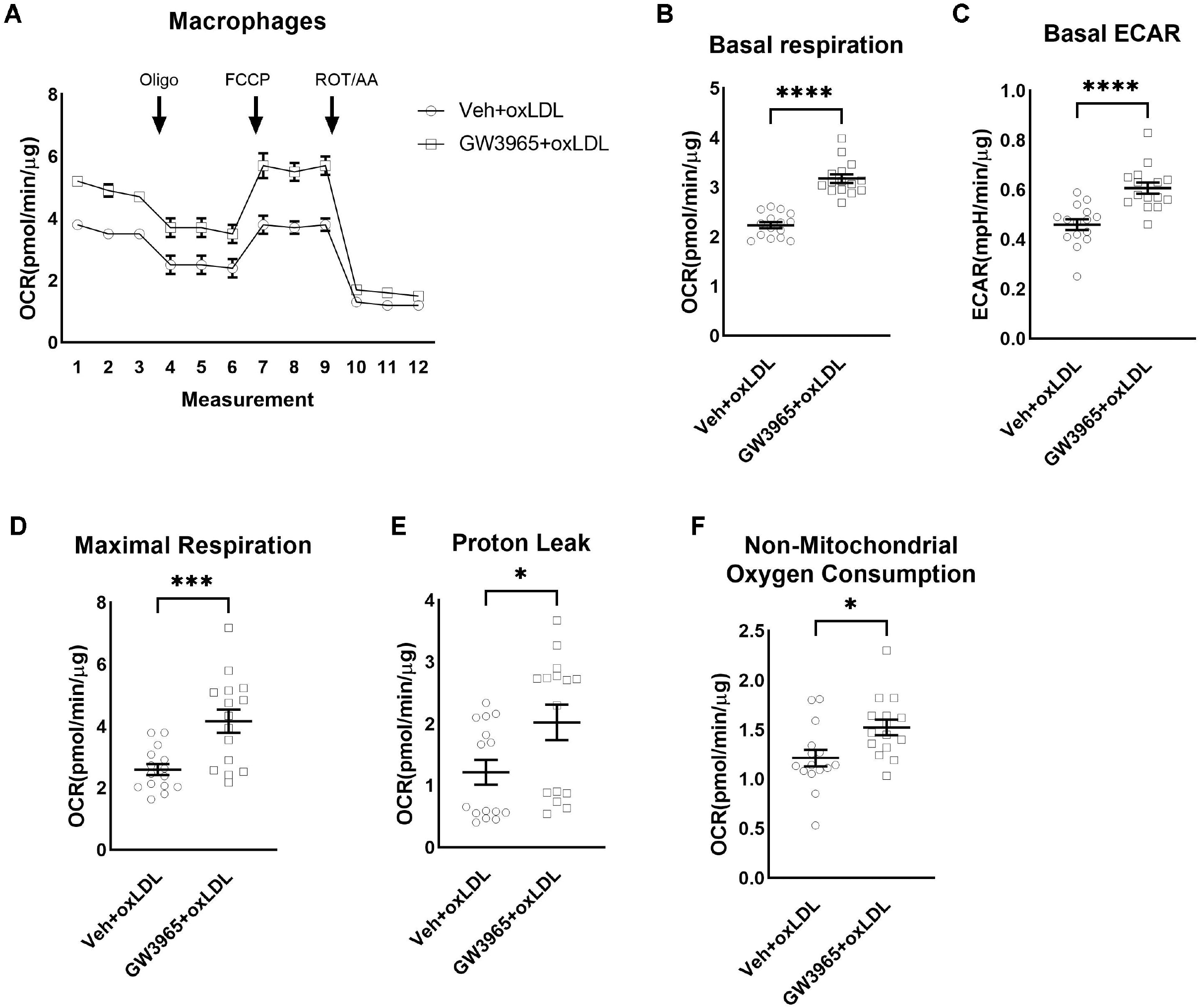
Activation of LXRs is associated with change in macrophage polarization. During oxygen consumption assay in the presence of oxldl and with and without LXRs agonist GW3965 (A), LXRs activation significantly increased basal respiration (B), basal ECAR (C), maximal respiration (D), proton leak (E), and non-mitochondrial oxygen consumption (F) (n=15 wells/group). All data are Mean±SEM. *p<0.05, ***p<0.0005, ****p<0.00005.

## Discussion

Aging is a major risk for damage of the structure and function of the PNS [42]. We confirm previous studies showing that old mice (24 months old) exhibit sensory dysfunction, loss of distal fibers, elevated endoneurial macrophages. Many studies showed demyelination, remyelination and axonal lesions resulting in the increase in phagocyting macrophages; that if they are correctly cleared maintain healthy nerve system [14–17]. Previous literature using 24 months old mice showed that silencing of foamy macrophages in the PNS improved the nerve structure and function [7]. These data suggested that foamy macrophages are crucial in the development of age-related neuropathy; however, the mechanism is not clear. It is possible that the presence of phagocyting macrophages is adaptive and necessary for a healthy nerve structure and function; however, a dysfunction in the lipid efflux and clearance of phagocytic immune cells would lead to nerve damages.

We have previously reported that activation of LXR in the sensory neurons of the DRG is required for amelioration of obesity-induced allodynia, and that selective deletion of LXRs from sensory neurons enhances the neuropathic pain phenotype [9]. We therefore tested the hypothesis that activation of LXRs can be effective in delaying/improving the progression of and found that the LXR selective agonist GW3965 either reversed or delayed the progression of multiple indices of neuropathy. This is consistent with our prior studies that showed LXR agonist treatment prevents progression of obesity-induced allodynia by altering peripheral sensory neuron function and their interaction with associated cells [9, 43]. Our findings suggest that activation of LXRs may influence sensory perception of the aged mice via activation of LXRs in macrophages of the SN that induces the M2 phenotype involved in nerve regeneration. An exploratory study using two human sural nerves suggests an age-related difference in macrophage polarization and encourages future additional studies using larger numbers.

LXRs are oxidized cholesterol derivative ligand-activated transcription factors and consist of two isoforms LXRα and LXRβ [9, 24, 25, 43]. LXRs play an important role in cholesterol metabolism [26, 27]. Several studies have reported that activation of LXRs inhibit the development of atherosclerosis, a property attributed to LXR-mediated ABCA1 expression and cholesterol efflux in macrophages [44, 45]. LXRs also play an important role in the regulation of cytokine production and the anti-inflammatory response [26, 27]. In this study, we provide evidence supporting the role of LXR in the regulation of a microenvironment permissive for neuronal regeneration. A number of anti-inflammatory mechanisms have been proposed to explain the actions of LXRs in other tissues including direct repression of pro-inflammatory gene promoters, cholesterol efflux, changes in plasma membrane signaling systems via modulation of membrane lipid composition and increased synthesis of fatty acids with anti-inflammatory activity [26, 46, 47]. The engulfment of myelin and other cellular debris in injured nerve by macrophages is associated with acute changes in their cellular lipid levels [29], including intracellular-free cholesterol. Activation of LXRs during this process promotes the efflux of free cholesterol by activation of ABCA1 [44, 45].

Our data demonstrate that cholesterol/lipid loaded macrophages are present in the SN of the aged mice and that activation of LXR decreases this lipid load in vivo and in vitro. There is novel body of evidence indicating that cellular metabolism determine macrophage function as either pro- or anti-inflammatory. However, the role of lipid content in PNS macrophage polarization is unknown. Assessing the bioenergetic profiles of macrophages treated with oxLDL and with and without LXRs activation, we find that macrophages treated with oxLDL have lower OCR and ECAR while macrophages treated with oxLDL and GW3965 have higher OCR and ECAR (highly energetic). Our metabolic flux analyses showed that both glycolysis and mitochondrial respiration are upregulated in macrophages treated with oxLDL and GW3965, suggesting that they could have a M2 phenotype [48]. Altogether, our data indicate that activation of LXRs in the macrophages of the PNS pathological microenvironment could induce phenotypic alteration by restoring or activating the lipid efflux in peripheral macrophages and that, LXRs may delay progression of neuropathic pain in aged mice.

## Author contributions

CKG and VMA were involved in the conception and design of the experiments. CKG, NE, AVG, GA, NAC, and VMA were involved in data collection, assembly, analysis and interpretation of data. CKG and VMA drafted the manuscript. CKG, NE, AVG, GA, NAC, and VMA revised the manuscript.

## Acknowledgements

We thank Loyola University Chicago animal facility for mice housing. This work was supported by NIH R01 DK117404 to VMA.

